# Afucosylated immunoglobulin G responses are a hallmark of enveloped virus infections and show an exacerbated phenotype in COVID-19

**DOI:** 10.1101/2020.05.18.099507

**Authors:** Mads Delbo Larsen, Erik L. de Graaf, Myrthe E. Sonneveld, H. Rosina Plomp, Federica Linty, Remco Visser, Maximilian Brinkhaus, Tonći Šuštić, Steven W. de Taeye, Arthur E.H. Bentlage, Jan Nouta, Suvi Natunen, Carolien A. M. Koeleman, Susanna Sainio, Neeltje A. Kootstra, Philip J.M. Brouwer, Rogier W. Sanders, Marit J. van Gils, Sanne de Bruin, Alexander P.J. Vlaar, Amsterdam UMC COVID-19 biobank study group, Hans L. Zaaijer, Manfred Wuhrer, C. Ellen van der Schoot, Gestur Vidarsson

## Abstract

IgG antibodies are crucial for protection against invading pathogens. A highly conserved N-linked glycan within the IgG-Fc-tail, essential for IgG function, shows variable composition in humans. Afucosylated IgG variants are already used in anti-cancer therapeutic antibodies for their elevated binding and killing activity through Fc receptors (FcγRIIIa). Here, we report that afucosylated IgG which are of minor abundance in humans (∼6% of total IgG) are specifically formed against surface epitopes of enveloped viruses after natural infections or immunization with attenuated viruses, while protein subunit immunization does not elicit this low fucose response. This can give beneficial strong responses, but can also go awry, resulting in a cytokine-storm and immune-mediated pathologies. In the case of COVID-19, the critically ill show aggravated afucosylated-IgG responses against the viral spike protein. In contrast, those clearing the infection unaided show higher fucosylation levels of the anti-spike protein IgG. Our findings indicate antibody glycosylation as a potential factor in inflammation and protection in enveloped virus infections including COVID-19.

## Main Text

Antibodies have long been considered functionally static, mostly determined by their isotype and subclass. The presence of a conserved N-linked glycan at position 297, in the so called constant Fc-domain of IgG, is essential for effector functions (*1*–*3*). Moreover, it is now generally accepted that the composition of this glycan is highly variable and has functional consequences (*2*–*4*). This is especially true for the core fucose attached to the Fc glycan. The discovery that IgG variants without core fucosylation cause elevated antibody dependent cellular cytotoxicity (ADCC), via increased IgG-Fc-receptor IIIa (FcγRIIIa) affinity (*5, 6*), resulted in next-generation glyco-engineered monoclonal antibodies (mAb) without core fucosylation for targeting tumors (*7*).

Generally, changes in the Fc glycans are associated with age, sex and autoimmune diseases (*8*). Serum IgG are highly fucosylated at birth and slightly decrease to ∼94% fucosylation at adulthood (*9*). Until now, no strong clues on how IgG core fucosylation is controlled have come forward.

We have previously observed that alloantibodies against red blood cells (RBC) and platelets show remarkably low IgG-Fc-fucosylation in most patients, even down to 10% in several cases (*10*–*12*), whereas the overall serum IgG Fc-fucosylation show consistently normal high levels. Moreover, we have reported the lowered IgG-Fc fucosylation to be one of the factors determining disease severity in pregnancy associated alloimmunizations, resulting in excessive thrombocytopenia’s and blood cell destruction when targeted by afucosylated antibodies (*11*–*13*). In addition to the specific afucosylated-IgG response against platelets and RBC antigens, this response has also been identified against HIV and Dengue virus (*14, 15*), but not for any other immune response so far, e.g. not against inactivated influenza, pneumococcal, meningococcal or tetanus vaccines (*16, 17*). Interestingly, low core fucosylation of anti-HIV antibodies has been suggested to be a feature of elite controllers of infection, and for Dengue it has been associated with enhanced pathology due to excessive FcγRIIIa-activation (*14, 15*).

Inspired by the similarities between the unique afucosylated IgG responses in various alloimmune responses (*10*–*12, 18*), HIV (*15*) and Dengue(*14*) – all being directed against surface exposed and membrane embedded proteins -we analyzed IgG glycosylation in anti-human platelet and red-blood cell alloimmune responses as well as in natural infections by other enveloped viruses, including HIV, cytomegalovirus (CMV), and SARS-CoV-2. Similarly, we also assessed for a non-enveloped virus (Parvovirus B19), vaccination with a protein subunit, and live attenuated enveloped viruses, to test if the antigen context is indeed an important determinant for IgG-Fc glycosylation.

To investigate the Fc-glycosylation of total- and antigen-specific antibodies, first IgG from >400 human serum samples was affinity-purified using protein G affinity beads and immobilized antigens, respectively. Thereafter, isolated IgG was digested with trypsin and resulting IgG1-Fc-glycopeptides were analyzed with liquid chromatography-mass spectrometry (LC-MS) (Fig. 2A) (*11, 16, 18*). Subsequently, intensities were extracted and IgG-glycosylation profiles were calculated (Fig. 1B-C).

**Fig. 1.**
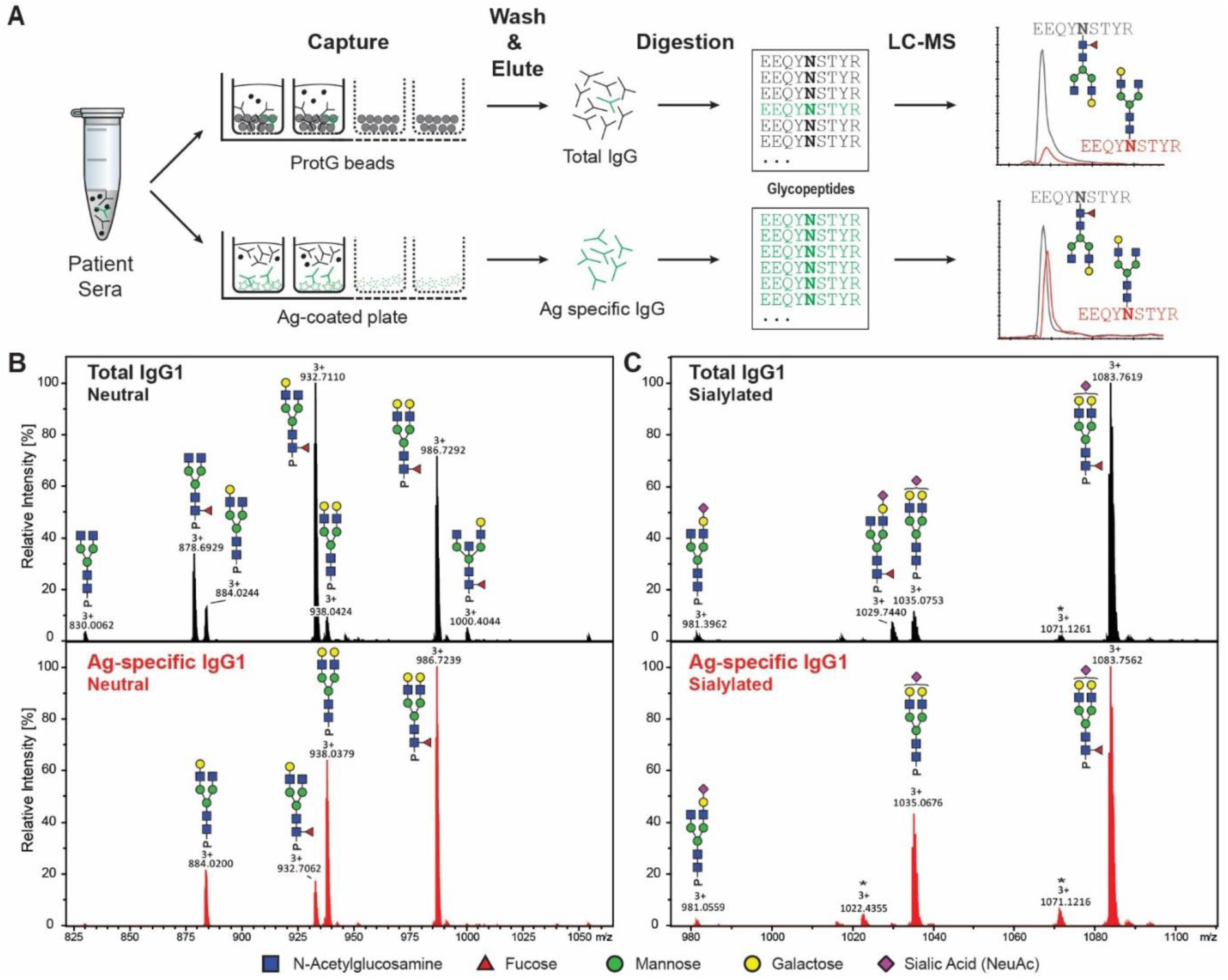
Flowchart of antibody specific IgG1 glycosylation analysis and mass spectrometric analysis. **A)** Antibodies from sera were captured using ProteinG beads and antigen-coated 96-well plates resulting in total and antigen-specific IgG fractions, respectively. Thereafter, isolated IgGs were digested with trypsin and the resulting glycopeptides were analyzed by nano liquid chromatography-coupled mass spectrometry. **B**,**C)** Representative mass spectra of glycopeptides encompassing the Fc glycosylation site Asn297. Neutral (**B**) and sialylated (**C**) IgG1 glycopeptides are shown from a single patients’ total (upper panel, in black) and antigen-specific (lower panel, in red) IgG1 fraction. Asterisks indicate non-Fc glycopeptides.

**Fig. 2.**
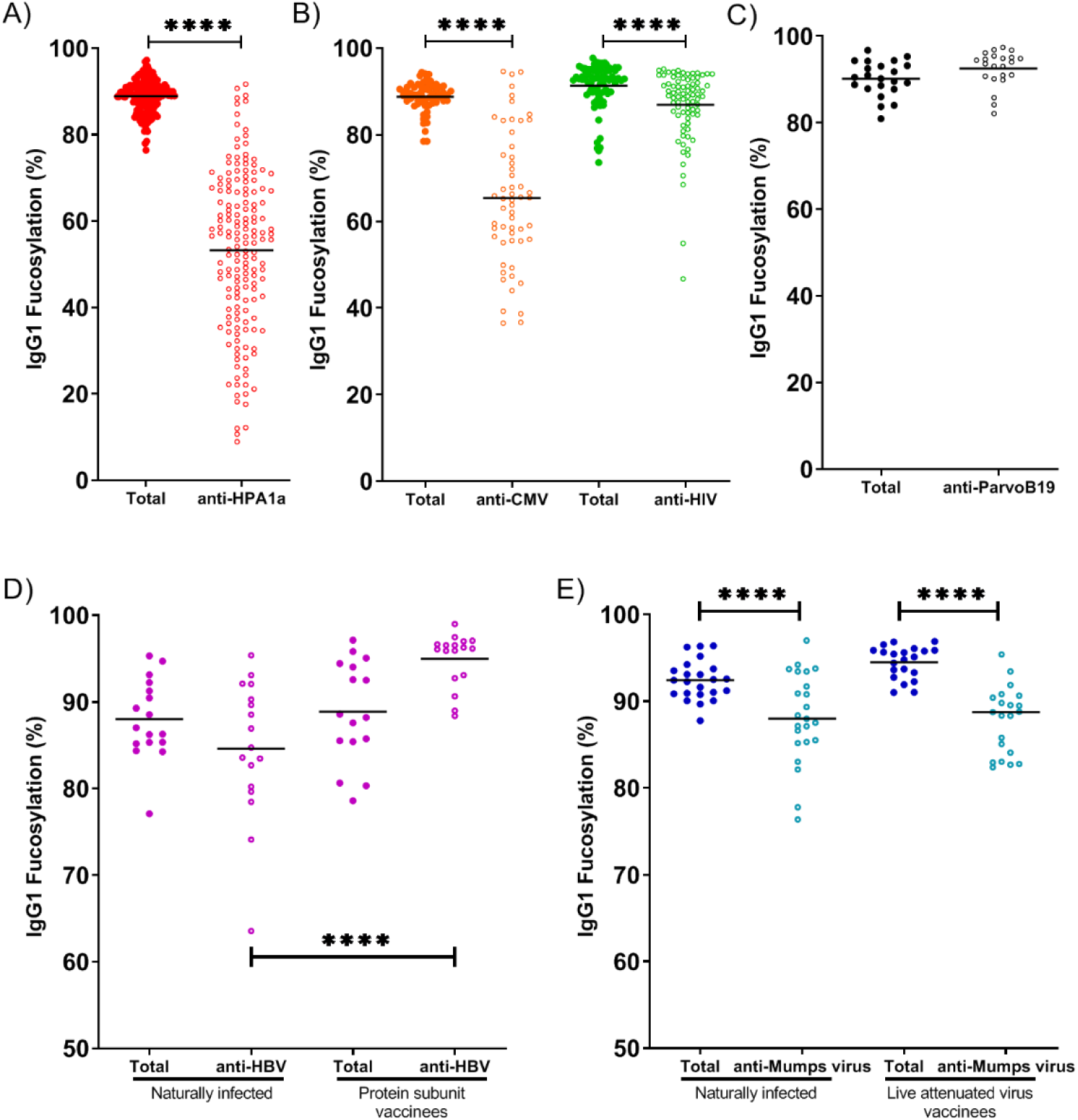
Foreign membrane protein antigens, such as envelope proteins of (attenuated) enveloped viruses or alloantigens can trigger afucosylated IgG responses. IgG1-Fc Fucosylation levels of total (filled circles) and antigen-specific (open circles) antibodies are shown for each differently color-coded group of antigens: **A)** alloantigen HPA-1a; **B)** viral envelope antigens from CMV and HIV; **C)** non-enveloped viral antigens from Parvo B19. **D)** IgG Fc-fucosylation levels of total and antigen-specific IgGs in individuals naturally infected with Hepatitis B Virus (left) or vaccinated with recombinant soluble HBsAg (right). **E)** Fc-fucosylation levels of total and ag-specific IgGs in individuals naturally infected with mumps virus (left), or vaccinated with live attenuated mumps virus (left). Statistical analysis was performed as paired t-test for A,B, and C and as a one-way ANOVA Sidak’s multiple comparisons test comparing total IgG to antigen-specific IgG within groups, and same specificity IgG between groups, for D and E. Only statistically significant differences are shown. *= p<0.1, **= p<0.01, ***= p<0.001, ****= p<0.0001.

Antigen-specific antibodies against the alloantigen human platelet antigen (HPA)-1a showed a strong decrease in fucosylation (Fig. 2A), similar to our previous findings for other alloantigens (*11, 18*).

Analogous to the platelet and Red Blood cell alloantigens (*10*–*12, 18*), the response to these enveloped viruses also showed significant afucosylation of the antigen-specific IgG (Fig.2B), while the afucosylation was absent against the non-enveloped virus Parvo B19 (Fig.2C). Of note, total IgG showed high fucosylation levels throughout (Fig.2A-C), underlining that the majority of human IgG responses consists of fucosylated IgG responses (*11, 16, 19*). The extent of the response to the enveloped viruses was highly variable, both between individuals and between the types of antigen, which is in agreement with the variable tendency of different RBC-alloantigens to induce an afucosylated response (*18*). Afucosylation was particularly strong for CMV and to a lesser degree for HIV (Fig. 2B). The anti-HIV response is in line with what was previously described by Ackerman et al., showing a decreased fucosylation of HIV-specific IgG compared to total IgG (*15*). Other glycan traits are depicted in Fig. S1.

To test whether some individuals had a greater intrinsic capacity to generate an afucosylated IgG response than others, we compared IgG-Fc fucosylation levels formed against different antigens within the same individual. No correlation in IgG1 Fc fucosylation was observed between anti-HPA1a and anti-CMV (Fig. S2), nor between anti-HIV and anti-CMV antibodies in the same individual (Fig. S2), suggesting that the level of afucosylation is not determined by a general host factor such as genetics but is rather stochastic or multifactorial, with the specific triggers remaining obscure.

To further investigate the immunological context by which potent low fucosylated IgG is formed, we compared immune responses to identical viral antigens in different contexts. First, we compared Hepatitis B surface antigen (HBsAg)-specific antibody glycosylation in humans naturally infected with Hepatitis B Virus (HBV) or vaccinated with the recombinant HBsAg protein (Fig. 2D). Whereas total-IgG1 fucosylation levels were similar in the two groups, anti-HBsAg IgG1 fucosylation was elevated in individuals vaccinated with the HBsAg protein both compared to the fucosylation of total IgG of both groups and antigen-specific IgG of the naturally infected group. The finding that HBsAg-specific antibodies in individuals that cleared a natural infection showed lowered Fc-fucosylation compared to protein subunit vaccination strongly suggests that a specific context for the antigenic stimulus is required for afucosylated-IgG responses.

We then compared antiviral-IgG responses against Mumps- and Measle viruses-formed after a natural infection or vaccination with live attenuated viruses. Unlike the HBV protein subunit vaccine, both attenuated live vaccines showed a similar Ag-specific Fc-fucosylation compared to their natural infection counterpart (Fig. 2E, Fig. S3). Both showed reduction, with a more prominent difference for the mumps response (Fig. 2E, Fig S3). Other glycan traits for anti-measles and anti-mumps are shown in Fig. S1.

We then tested if this type of response also plays a role in patients with SARS-CoV-2 (COVID-19). Symptoms of COVID-19 are highly diverse, ranging from asymptomatic or mild self-limiting infection to a severe airway inflammation leading to respiratory distress, often with a fatal outcome(*20, 21*). Both extreme trajectories, follow similar initial responses: patients have approximately a week of relatively mild symptoms, followed by a second wave that either clears the disease or leads to a highly-aggravated life-threatening phenotype (*20, 21*). Both the timing of either response type and the differential clinical outcome suggests different paths taken by the immune system to combat the disease. So far no clear evidence has emerged that can make a distinction between these two hypothetical immunological paths. In accordance with our theory and responses seen in other enveloped viruses, anti-S IgG responses against SARS-CoV-2 spike protein (S), expressed on cell surface/viral envelope, were strongly skewed towards low core fucosylation, while those against the nucleocapsid protein (N), not expressed on cell surface/viral envelope, was characterized with high level of fucose. Importantly, the afucosylated anti-S IgG-responses of patients with Acute Respiratory Distress Syndrome (ARDS) hospitalized in intensive care units were significantly lower than in convalescent plasma donors consisting of individuals who were asymptomatic or had relative mild symptoms (non-ARDS) (Fig. 3A).

**Figure 3:**
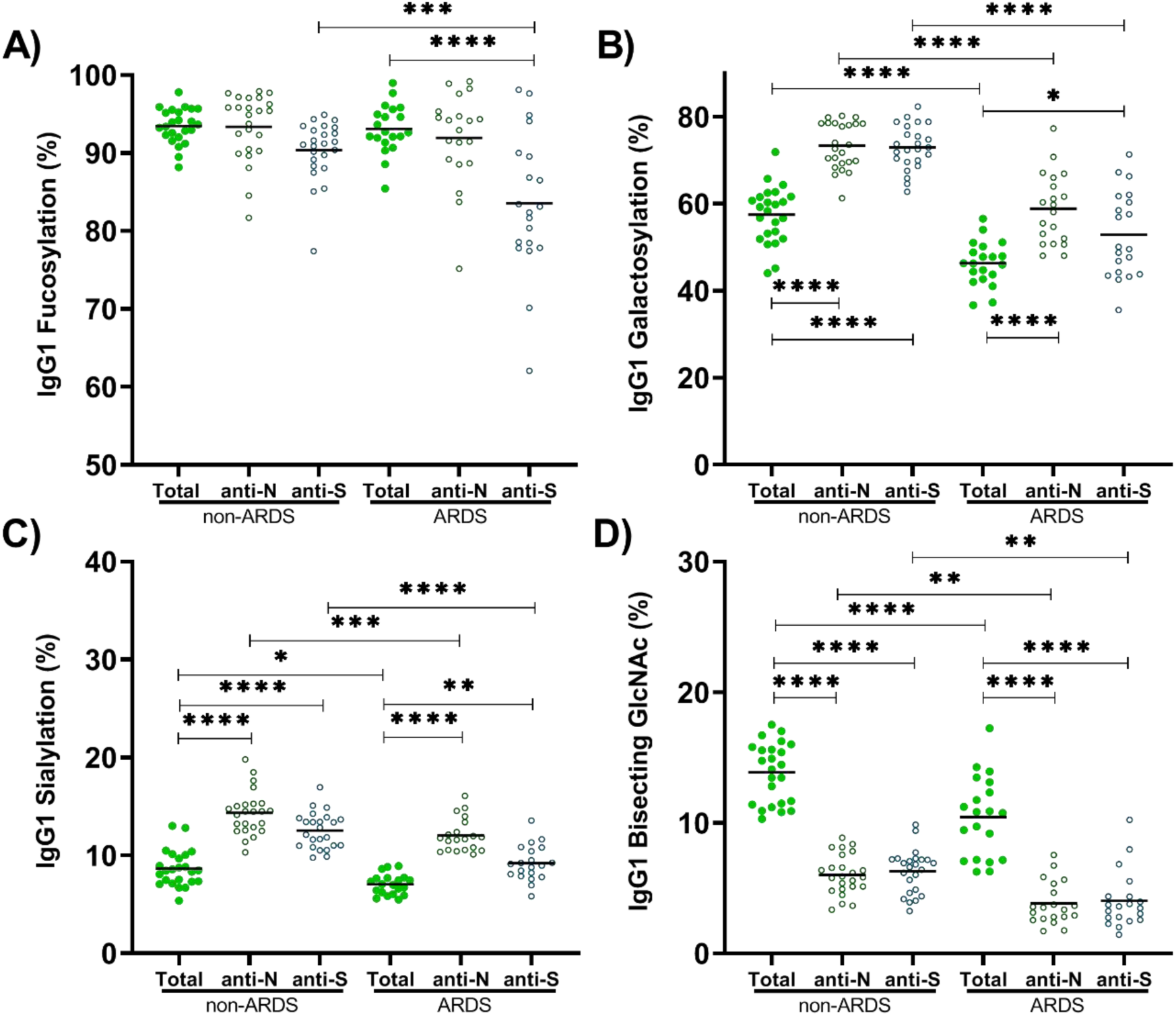
Fucosylation levels of anti-S IgG1, but not anti-N, are significantly decreased in critically ill COVID-19 patients. **A)** Fucosylation was significantly lowered for ARDS patients for anti-S, but not in non-ARDS donors naturally clearing the infection. Anti-N responses were similar to total IgG. **B)** Galactosylation, was increased for both anti-N and anti-S in non-ARDS donors as well as ARDS patients **C)** Sialylation shows similar pattens as galactosylation, and **D)** Bisection was very low for both anti-N and anti-S in both patient groups. Statistical analysis was performed as a one-way ANOVA with Sidak’s multiple comparisons test, comparing total IgG to antigen-specific IgG within groups, and same specificity IgG between groups. Only statistically significant differences are shown. *= p<0.1, **= p<0.01, ***= p<0.001, ****= p<0.0001

This lowered fucosylation of the anti-S was not a general issue of the inflammation as total IgG-fucosylation levels were similar in the two groups and to what has been reported in the general population (∼94%) (*11, 16*). In addition, IgG galactosylation of both anti-S and anti-N responses tended to be higher than seen in total IgG, compatible with increased galactosylation observed in active or recent immunization (*16, 22*). However, IgG galactosylation levels in general were lower in the ARDS patients, perhaps suggesting lower capacity to clear the infection by reduced complement activity (*23*). This may be a reflection of a slight age difference in these two groups (non-ARDS donors median 53±12, ARDS patients 61±7.9), as galactose generally decreases slightly with advancing age (*9, 19*). More importantly, the lowered fucosylation in the anti-S responses of the ARDS patients, strongly suggest a pathological role through FcγRIIIa, similar to what was previously proposed for Dengue (*14*). In Dengue, non-neutralizing antibodies formed to previous infections of other Dengue serotypes, also tend to have low level of core-fucosylated IgG and, as they are not capable of preventing infection, leading to aggravated Dengue-disease due to FcγRIIIa-mediated overreactions by immune cells (*14*).

In conclusion, our results show a pattern of afucosylated IgG1 immune responses against membrane-embedded antigens such as surface membrane proteins of allo-antigens on blood cells or on enveloped viruses, in contrast to soluble protein antigens and non-enveloped viruses for which immune responses with high levels of IgG1 fucosylation are consistently observed. We hypothesize that antigen-presenting membranes are directly sensed by B cells combining at least two signals provided by the B cell receptor and a yet unknown host receptor-ligand pair, not occurring for soluble proteins, internal protein of enveloped viruses or non-enveloped viruses(Fig. 4).

**Fig. 4.**
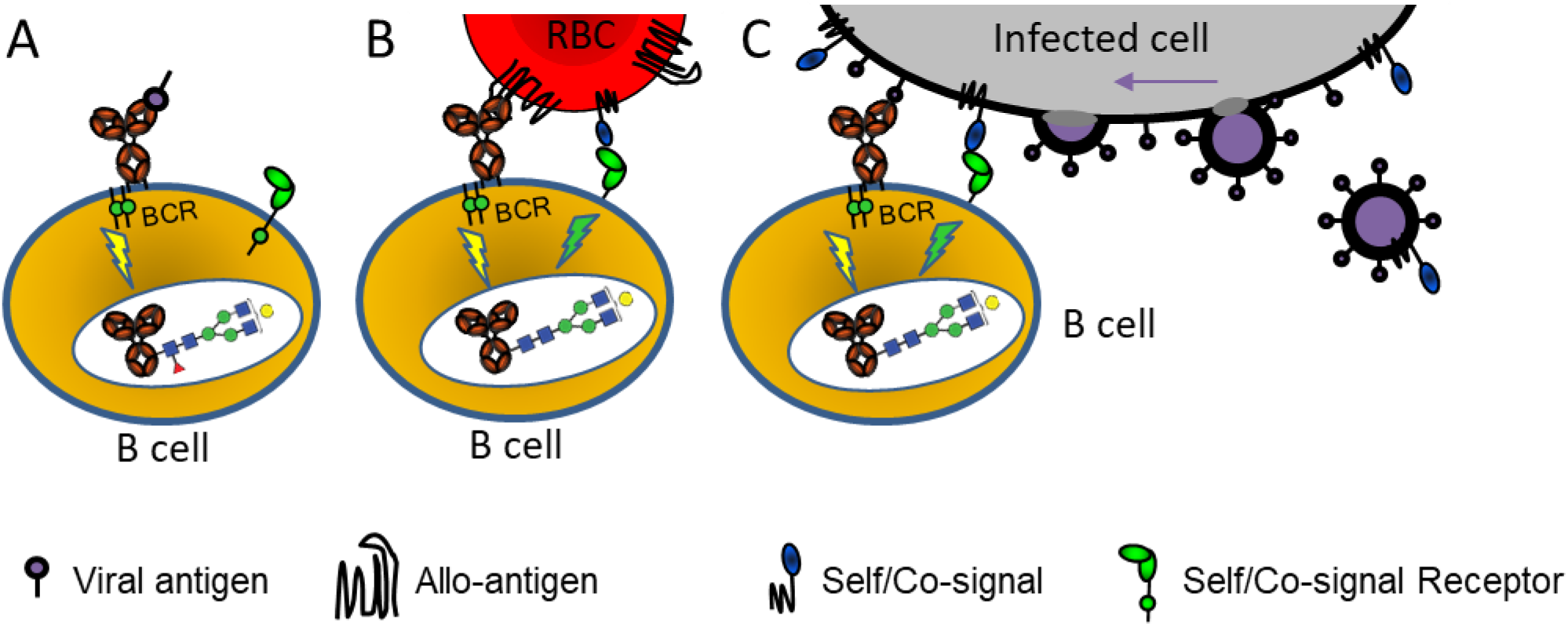
Hypothetical model explaining how the context of antigen can lead to altered immune signaling, giving rise to altered IgG-glycosylation. **A)** Immune response to soluble protein antigen: B-cell receptor (BCR, a membrane immunoglobulin) is activated, resulting in the production of normal fucosylated antibodies. **B)** Immune response to allo-antigens: Paternal allo-antigen on a Red Blood Cell (RBC) recognized by the BCR, and possibly by a yet unknown immune regulatory receptor-ligand pair providing a signal for self. **C)** Immune response to enveloped viral infection: Recognition of enveloped-virus infected cells by B cells is similar as for cellular-alloantigen recognition. The initial recognition can potentially occur towards enveloped-virally infected cells and possibly after viral assembly (far right). The proposed signaling in b-c) causes altered glyco-programming of the B cells culminating in a unique IgG-response characterized by a low fucosylation (fucose red triangle) and enhanced ADCC. The model potentially explains both why immune response to soluble proteins, non-enveloped viruses and cellular pathogens such as bacteria is different to responses to enveloped-viruses and why immune responses to allo-antigens mimic that of an enveloped viral infection.

Alternatively, the differential recognition may be more complex and require additional interactions from antigen-presenting cells, T cells and/or cytokines. Importantly, our studies imply that a membrane context may be necessary but not always sufficient to trigger an immune response with high levels of afucosylated IgG. (*18*). This translates into a vast spread of afucosylation levels between individuals as well as for distinct responses of the same individual against different antigens. The large difference in the level of antigen-specific afucosylated responses observed between patients contributes to the variability of disease severity, as has been shown for the neonatal alloimmune cytopenias (*11, 12, 18*), Dengue (*14*) and now also for COVID-19. This underscores the significance for diagnosis of possible disease trajectories and guides future treatments aimed at minimizing this FcγRIIIa-stimulus.

Importantly, when IgG afucosylation does occur, the final outcome, results in a potent immune response, honed for destruction of targets cells by FcγRIIIa-expressing NK cells, monocytes and macrophages but also FcγRIIIb-expressing granulocytes. This can be desired in some responses such as against HIV (*15*), which can be achieved with available attenuated enveloped viral vaccine shuttles (*24*) against difficult targets. On the other hand, this can also lead to an undesired exaggerated response, as is apparent for both Dengue (*14*) and COVID-19. This is exemplified in experiments in monkeys, where vaccination with Modified Vaccinia Ankara virus ferrying spike proteins of SARS-CoV lead to strong ADE response (*25*) mimicking pathologies in critically ill SARS-CoV2 patients (*21*). This suggests that a subunit protein vaccine to be a safer option as seen in rat models for SARS-CoV2 (*26*), unless the vaccine also induces a strong neutralizing effect.

For COVID-19, the data suggest that afucosylation of anti-S IgG may contribute to the exacerbation of the disease in an subset of patient ending up in Intensive care units with ARDS. Thus although they can be protective, they might potentially behave as double-edged swords, and may contribute to the observed cytokine storm (*27*). As such this has direct consequences for improving current therapies with IVIg, convalescent plasma and the route taken towards vaccine development. In addition, the suggested role of afucosylated antibodies in the pathogenesis of COVID-19 might open new opportunities for therapy of this disease. Future attempts of generating high-titer COVID-19 immunoglobulin treatments, should preferably use plasma enriched in fucosylated anti-COVID-19 antibodies. These may outcompete afucosylated anti-SARS-CoV2 IgG-responses developing in the patients to avoid symptom escalation and promote virus neutralization.

## Materials and Methods

### Patient samples

Healthy blood donor samples from Sanquin, Amsterdam, The Netherlands, were used to analyze ParvoB19 (n=22), Measle virus (n=21 natural infection, n=24 Live Attenuated vaccine), Mump virus (n=21 natural infection, n=24 Live Attenuated vaccine) and HBV antibodies (n=17 natural infection, n=16 HBsAg vaccination). Anti-HPA-1a samples have been described elsewhere (*13*). HIV-samples (n=80) from the Amsterdam Cohort Studies on HIV infection and AIDS (ACS) were used to analyze HIV-specific antibody glycosylation. SARS-CoV2 patient samples from ICU patient from the Amsterdam UMC COVID study group were included, as well as from Sanquin blood donors found positive. The ACS have been conducted in accordance with the ethical principles set out in the declaration of Helsinki and all participants provided written informed consent. The study was approved by the Academic Medical Center institutional Medical Ethics Committee of the University of Amsterdam.

Peripheral blood samples from patients with HPA-1a alloantibodies and CMV-specific antibodies (n=62) were collected by the Finnish Red Cross Blood service, Platelet Immunology Laboratory, Helsinki, Finland.

### Purification of CMV-specific antibodies from sera

CMV-specific antibodies were purified using antigen-coated plates (Serion ELISA classic, Cytomegalovirus IgG, Würzburg, Germany). Sera (20 *µ*L) diluted in specimen diluent (80 *µ*L) from kit was incubated for 1 hour at 37 °C degrees. Positive and negative controls from the kit and CMV-negative patients samples were used as controls. The plates were washed seven times: three times with 300 *µ*L wash-buffer from the kit, followed by two washed with the same volume Phosphate buffered saline (PBS) and deionized water. The bound antibodies were then eluted using 100 *µ*L of 100 mmol/l formic acid. No IgG was found in eluates from blank wells and CMV-negative patients samples.

### Purification of Measle- and Mump-virus specific antibodies from sera

Ag-specific antibodies were purified using antigen-coated plates (Serion ELISA classic, Measles IgG and Mumps IgG, Würzburg, Germany). Sera (20 *µ*L) diluted in specimen diluent (80 *µ*L) from kit was incubated for 1 hour at 37 °C degrees. Positive and negative controls from the kit were used as controls. The plates were washed seven times: three times with 300 *µ*L wash-buffer from the kit, followed by two washed with the same volume Phosphate buffered saline (PBS) and 50mM ammonium bicarbonate. The bound antibodies were then eluted using 100 *µ*L of 100 mM formic acid. IgG was found in the eluates of positive controls and no IgG was found in eluates from blank wells and negative control samples.

### Purification of HBV-specific antibodies from sera

To isolate HBsAg specific antibodies from patients after infection and vaccination, HBs antigen-coated plates (ETI-AB-AUK-3, Diasorin, Schiphol-Rijk, The Netherlands) were used. Sera were diluted five times in specimen diluent from kit (20 *µ*L serum with 80 *µ*L diluent) and incubated for 1 hour at room temperature with shaking 450 r.p.m. (Heidolph Titramax 100, Schwabach, Germany). HBV-naive and HBV-resolved samples from Sanquin, Amsterdam, The Netherlands were used as controls. Washing and eluting specific antibodies was done as described above for CMV-specific antibodies.

### Purification of HIV-specific antibodies from sera

HIV-specific antibodies were isolated using HIV antigen-coated plates (Murex HIV1.2.0 kit 9E25-01, Diasorin, Schiphol-Rijk, The Netherlands). Sera were diluted two times in sample diluent from kit (50 *µ*L serum with 50 *µ*L diluent) and incubated for 1 hour at room temperature shaking 450 r.p.m. (Heidolph Titramax 100, Schwabach, Germany). As positive control, anti-HIV gp120 monoclonal was used (IgG1 b12; 100 *µ*g purified antibody in PBS at 1 mg/ml; NIH Aids Reagent Program, La Jolla, CA, US). Washing and eluting specific antibodies was done as described above for CMV-specific antibodies.

### Purification of ParvoB19-specific antibodies from sera

ParvoB19-specific antibodies were isolated using ParvoB19 antigen-coated plates (Abcam1788650-Anti-Parvovirus B19 IgG ELISA, Cambridge, United Kingdom). Sera were diluted five times in sample diluent from kit (20 *µ*L serum with 80 *µ*L diluent) and incubated for 1 hour at room temperature shaking 450 r.p.m. (Heidolph Titramax 100). Positive and negative controls from the kit were used as controls. Washing and eluting specific antibodies was done as described above for CMV-specific antibodies.

### Purification of anti-N and anti-S specific antibodies from plasma

SARS-Cov-2-specific antibodies were purified using antigen-coated plates (NUCN, Roskilde, Denmark). Plates were coated (over-night, 4°C) with recombinant trimerized spike protein produced as described recently (*28*) or N protein (accession number MN908947, produced in HEK cells with HAVT20 leader peptide, 10xhis tag and a Brit tag as in (*23*)) in PBS(5 µg/mL and 1 µg/mL, respectively). Plates were washed 3× with PBS supplemented with 0.05 % TWEEN 20^®^ (PBS-T) and plasma (20 *µ*L) diluted in PBS-T (180 *µ*L) was incubated on the plates(1 hour, 37 °C, shaking). Sera dating pre COVID-19 pandemic were used as negative controls. The plates were washed seven times: 3× with PBS-T, 2× with PBS and 2× with ammonium bicarbonate (50mM). The bound antibodies were then eluted with formic acid (100 mM, 5 min, shaking).

### Purification of total IgG from sera

Total IgG1 antibodies were captured from 2 *µ*L of serum using Protein G Sepharose 4 Fast Flow beads (GE Healthcare, Uppsala, Sweden) in a 96-well filter plate (Millipore Multiscreen, Amsterdam, The Netherlands) as described previously (*11*) or by using Protein G cartridges on the AssayMAP Bravo (Agilent Technologies, Santa Clara, USA) Briefly, 1 µL serum diluted in PBS were applied to the cartridges, followed by washes of PBS, LC-MS pure water and finally eluted with formic acid (1%).

### Mass spectrometric IgG-Fc glycosylation analysis

Eluates containing either antigen-specific antibodies or total IgG were collected in V-bottom plates, dried by vacuum centrifugation for 2.5 hours at 50°C. The HPA1a, CMV, HIV, ParvoB19, HBV, and COVID-19 samples were then subjected to proteolytic cleavage using trypsin as described before (*11*). The measles and mumps cohort samples were dissolved in a buffer containing 0.4% sodium deoxycholate(SDC), 10mM TCEP, 40mM chloroacetamide, 100 mM TRIS pH8.5. After 10min incubation at 95C, 250 ng Trypsin in 50 mM ammonium bicarbonate was added. The digestion was stopped after an overnight incubation by acidifying to 2% formic acid. Prior to MS injection, SDC precipitates were removed by centrifuging samples at 20 000 rcf for 30 minutes.

Analysis of IgG Fc-glycosylation was performed with nanoLC reverse phase (RP)-electrospray (ESI)-MS on an Ultimate 3000 RSLCnano system (Dionex/Thermo Scientific, Breda, The Netherlands) coupled to an amaZon speed ion trap MS (Bruker Daltonics, Bremen, Germany) (*11*) for all samples except for the Measles and mumps cohorts that were measured on a Impact HD quadrupole-time-of-flight MS (Bruker Daltonics). (Glyco-)peptides were trapped with 100% buffer A (0.1% formic acid in water) and separated on a 15 min 0-25% buffer B (95% acetonitrile, 5% water) linear gradient. In the current study we focused on IgG1, without analyzing IgG3 due to its possible interference with IgG2 and IgG4 at the glycopeptide level (*29*). Mass spectrometry results were extracted and evaluated using FlexAnalysis software (Bruker Daltonics) for all samples except for the Measles virus, and Mumps virus cohorts that were analyzed with Skyline software. Data was judged reliable when the sum of the signal intensities of all glycopeptide species (Table S1) was at least higher than background plus 10 times its standard division, otherwise the data was excluded (*11*). The total level of glycan traits was calculated as described in Table S2.

### Statistical analysis

Statistical analyses were performed using GraphPad Prism version 7.02 for Windows (GraphPad Software Inc., La Jolla, CA, www.graphpad.com). To analyze whether Fc-fucosylation for total and antigen specific IgG differs between the tested cohorts, statistical analysis was performed using t tests. To investigate whether Fc-fucosylation profiles of two specific antibodies in the same individual are correlated statistical analysis was performed using a Pearson correlation. The level of significance was set at P<0.05.

## Acknowledgments

We thank prof. dr. R.J.M. ten Berge for helpful discussions. The Amsterdam Cohort Studies on HIV infection and AIDS, is a collaboration between the Amsterdam Health Service, the Academic Medical Centre of the University of Amsterdam and Sanquin Blood Supply Foundation. The ACS is part of the Netherlands HIV Monitoring Foundation and financially supported by the Netherlands National Institute for Public Health and the Environment. We are greatly indebted to all cohort participants for their continuous participation.

## Author contributions

All samples were collected by SN, SS, ST, AV, AB, RV, MB, TS, SB, FL, NK, HZ, and MS. MS, CK, RP, JN, ML and EdG performed antibody purifications, mass spectrometric analyses and data processing. FL, MB, TS, and PB, generated recombinant antigens, MS, EdG, GV, and EvdS analyzed clinical data and performed data analysis. ML, MS, EdG, MW and GV made figures and tables. MG, RS, EvdS, MW and GV supervised the study. All authors contributed to analysis and interpretation of the data. ML, MS, EdG, ES, MW and GV wrote the paper, which was critically revised and approved by all authors.

## Data and materials availability

The data presented in this manuscript are in the main paper.

## Suplementary data

**Fig. S1:**
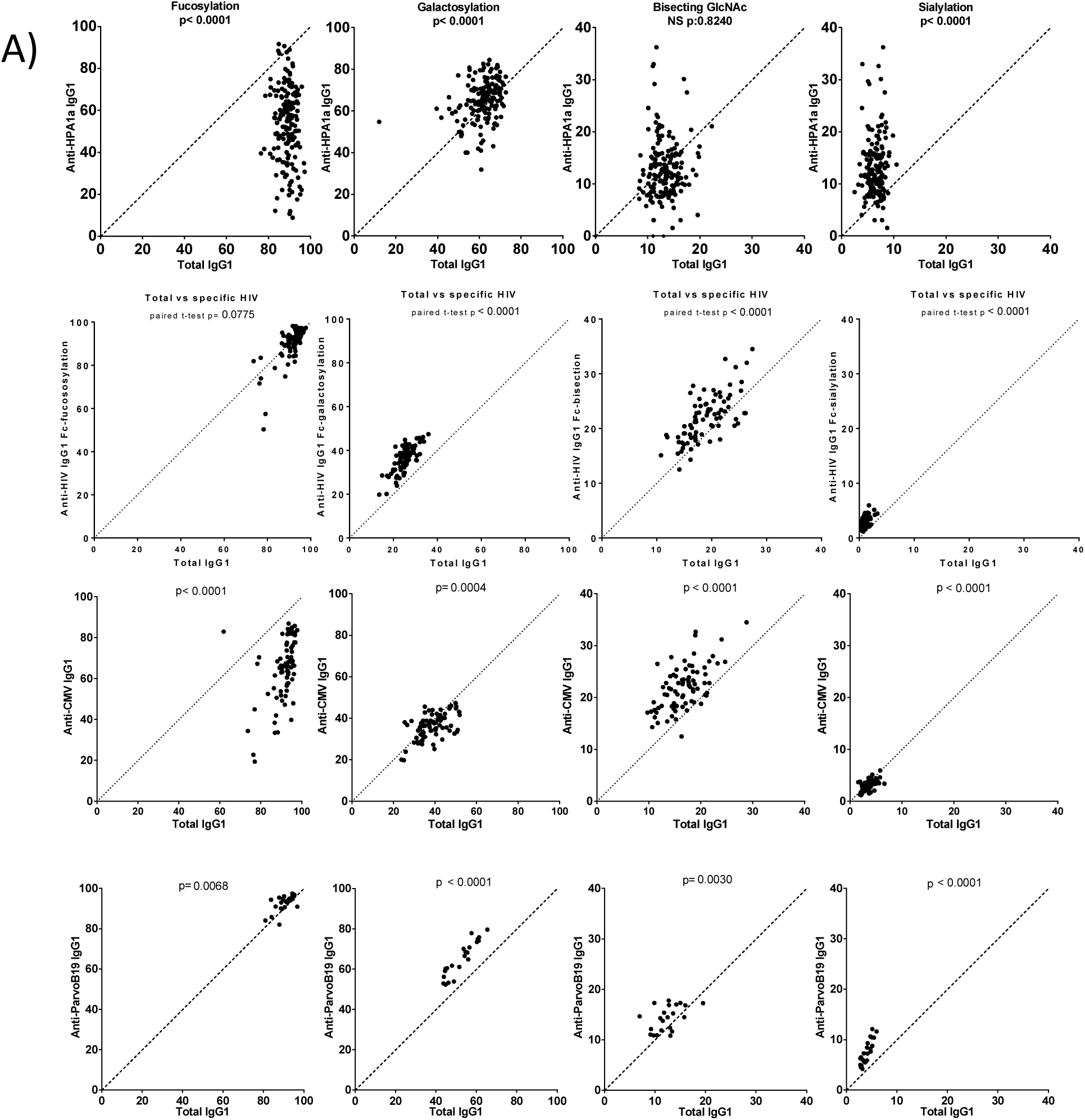
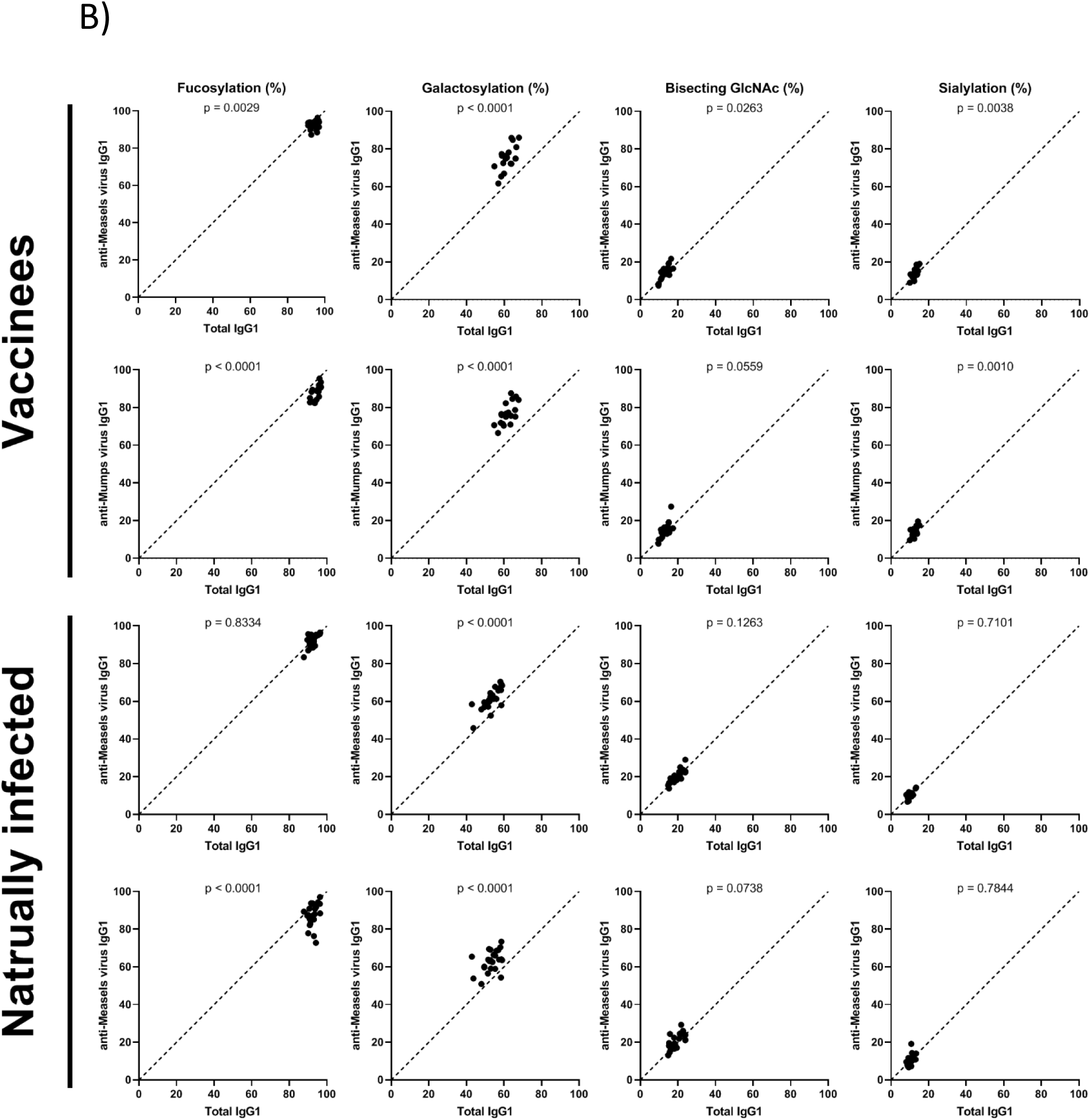
IgG1-glycosylation for A) anti-HPA-1a, HIV, CMV and Parvo-B19 and B) anti-Measles and Mumps (next page). Shown are total IgG glycan traits (x-axis) vs the corresponding antigen-specific glycosylation on the Y-Axis for Fucosylation, Galactosylation, Bisection and sialylation.

**Fig. S2.**
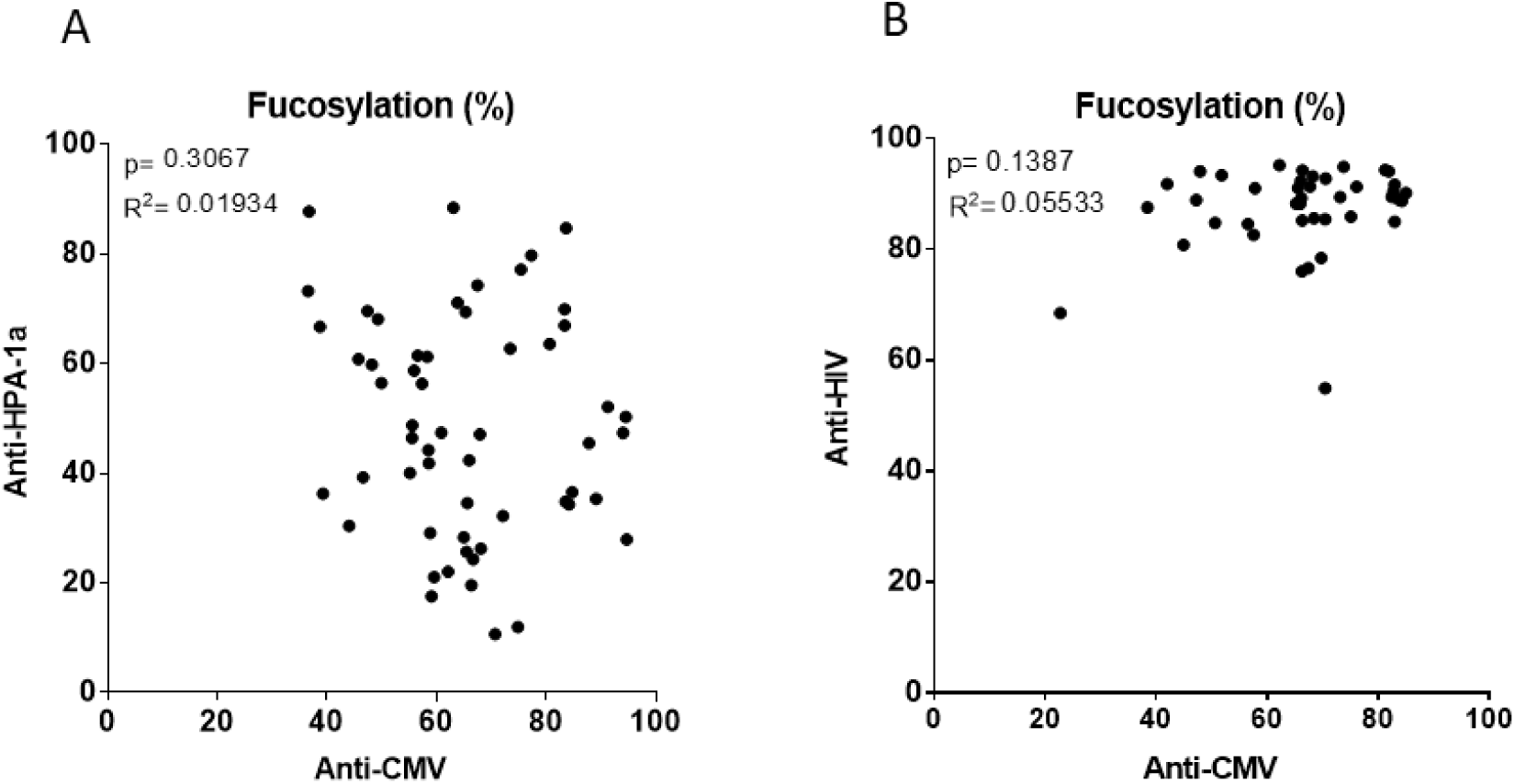
Antibody fucosylation is not determined by a general host factor. **a.** No correlation was found between the level of IgG1 Fc-fucosylation made during alloimmunization against HPA1a in pregnancy (y-axis) and CMV (x-axis) in the same individual. **b.** Also no correlation was found between the level of IgG1 Fc-fucosylation made against HIV (y-axis) and CMV (x-axis) in the same individual. Statistical analysis was performed using Pearson correlation.

**Fig. S3:**
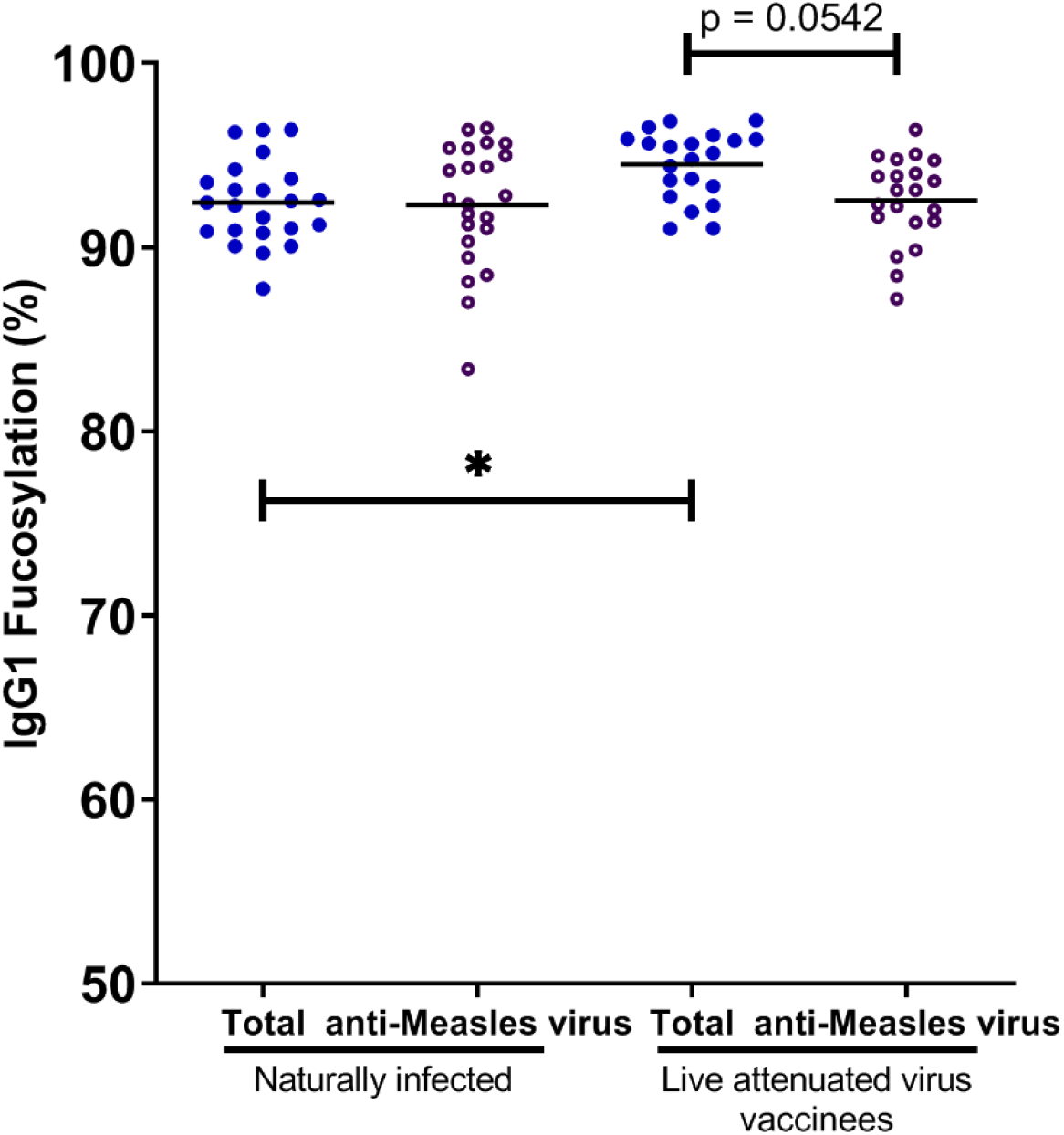
Fucosylation of anti-Measles. Compared to total IgG fucosylation, the antigen-specific IgG fucosylation, of anti-measle antibodies was only significantly lowered in the younger vaccinated cohort (mean age 19.5). This is likely to be masked by the natural tendency of lowered total IgG-fucosylation with increasing age (9), as the naturally infected cohort (before introduction of the MeV/MuV vaccination program in 1980s in the Netherlands) is older than the vaccine cohort (average 63.5 vs 19.5 years, respectively). In line with this, the total IgG fucosylation of the older cohort showed significantly lowered total-IgG fucosylation compared to the younger vaccinated cohort / Statistical analysis was performed as paired t-test for A,B, and C and as a one-way ANOVA Sidak’s multiple comparisons test comparing total IgG to antigen-specific IgG within groups, and same specificity IgG between groups, for D and E. Only statistically significant differences are shown. (*= p<0.1)

**Supplementary Table 1.** Overview of the IgG Fc glycopeptides which were included. The monoisotopic m/z value of the 2+ and 3+ charge state are shown, together with the average retention time determined for all N-glycans of each IgG subclass. Glycan compositions are denoted using the following nomenclature: H = hexose, N = N-acetylhexosamine; F = fucose; S = sialic (N-acetylneuraminic acids).

| N-glycopeptides | m/z 2+ | m/z 3+ | retention time (s) |
| --- | --- | --- | --- |
| IgG1 H3N4F1S0 [G0F] | 1317.526 | 878.687 | 336 |
| IgG1 H4N4F1S0 [G1F] | 1398.552 | 932.704 | 336 |
| IgG1 H5N4F1S0 [G2F] | 1479.579 | 986.722 | 336 |
| IgG1 H3N5F1S0 [G0FN] | 1419.066 | 946.380 | 336 |
| IgG1 H4N5F1S0 [G1FN] | 1500.092 | 1000.398 | 336 |
| IgG1 H5N5F1S0 [G2FN] | 1581.119 | 1054.415 | 336 |
| IgG1 H3N4F0S0 [G0] | 1244.497 | 830.001 | 336 |
| IgG1 H4N4F0S0 [G1] | 1325.524 | 884.018 | 336 |
| IgG1 H5N4F0S0 [G2] | 1406.550 | 938.036 | 336 |
| IgG1 H3N5F0S0 [G0N] | 1346.037 | 897.694 | 336 |
| IgG1 H4N5F0S0 [G1N] | 1427.063 | 951.712 | 336 |
| IgG1 H5N5F0S0 [G2N] | 1508.090 | 1005.729 | 336 |
| IgG1 H4N4F1S1 [G1FS] | 1544.100 | 1029.736 | 396 |
| IgG1 H5N4F1S1 [G2FS] | 1625.127 | 1083.754 | 396 |
| IgG1 H4N5F1S1 [G1FNS] | 1645.640 | 1097.429 | 396 |
| IgG1 H5N5F1S1 [G2FNS] | 1726.667 | 1151.447 | 396 |
| IgG1 H4N4F0S1 [G1S] | 1471.071 | 981.050 | 396 |
| IgG1 H5N4F0S1 [G2S] | 1552.098 | 1035.068 | 396 |
| IgG1 H4N5F0S1 [G1NS] | 1572.611 | 1048.743 | 396 |
| IgG1 H5N5F0S1 [G2NS] | 1653.638 | 1102.7610 | 396 |
| IgG1 H5N4F1S2 [G2FS2] | 1770.675 | 1180.786 | 396 |

**Supplementary Table 2.** An overview of the calculations for the derived glycosylation traits.

| Supplemental Table 2. An overview of the calculations for the derived glycosylation traits. |  |  |
| --- | --- | --- |
| Derived trait | Definition | Calculation |
| Fucosylation | % of N-glycans which carry a core fucose | $\frac{[H3N4F1S0 + H4N4F1S0 + H5N4F1S0 + H3N5F1S0 + H4N5F1S0 + H5N5F1S0 + H4N4F1S1 + H5N4F1S1 + H4N5F1S1 + H5N5F1S1 + H5N4F1S2]}{\text{Sum of all glyco species}}$ |
| Galactosylation | % of N-glycans which carry a galactose | $\frac{[(H5N4F1S0 + H5N5F1S0 + H5N4F0S0 + H5N5F0S0 + H5N4F1S1 + H5N5F1S1 + H5N4F0S1 + H5N5F0S1 + H5N4F1S2) + 0.5 * (H4N4F1S0 + H4N5F1S0 + H4N4F0S0 + H4N5F0S0 + H4N4F1S1 + H4N5F1S1 + H4N4F0S1 + H4N5F0S1)]}{\text{Sum of all glyco species}}$ |
| Sialylation | % of N-glycans which carry a N-acetylneuraminic (sialic) acid | $\frac{[H5N4F1S2 + 0.5 * (H4N4F1S1 + H5N4F1S1 + H4N5F1S1 + H5N5F1S1 + H4N4F0S1 + H5N4F0S1 + H4N5F0S1 + H5N5F0S1)]}{\text{Sum of all glyco species}}$ |
| Bisection | % of N-glycans which carry a bisecting N-acetylglucosamine | $\frac{[H3N5F1S0 + H4N5F1S0 + H5N5F1S0 + H3N5F0S0 + H4N5F0S0 + H5N5F0S0 + H4N5F1S1 + H5N5F1S1 + H4N5F0S1 + H5N5F0S1]}{\text{Sum of all glyco species}}$ |

